# Neuronal wiring receptors Dprs and DIPs are GPI anchored and this modification contributes to their cell surface organization

**DOI:** 10.1101/2023.03.02.530872

**Authors:** Meike Lobb-Rabe, Wioletta I. Nawrocka, Robert A. Carrillo, Engin Özkan

## Abstract

The *Drosophila* Dpr and DIP proteins belong to the immunoglobulin superfamily of cell surface proteins (CSPs). Their hetero- and homophilic interactions have been implicated in a variety of neuronal functions, including synaptic connectivity, cell survival, and axon fasciculation. However, the signaling pathways underlying these diverse functions are unknown. To gain insight into Dpr–DIP signaling, we sought to examine how these CSPs are associated with the membrane. Specifically, we asked whether Dprs and DIPs are integral membrane proteins or membrane anchored through the addition of glycosylphosphatidylinositol (GPI) linkage. We demonstrate that Dprs and DIPs are GPI anchored to the membrane of insect cells and validate these findings for some family members in vivo using *Drosophila* larvae, where GPI anchor cleavage results in loss of surface labeling. Additionally, we show that GPI cleavage abrogates aggregation of insect cells expressing cognate Dpr–DIP partners. To test if the GPI anchor affects Dpr–DIP localization, we replaced it with a transmembrane domain and observed perturbation of sub-cellular localization on motor neurons and muscles. These data suggest that membrane anchoring of Dprs and DIPs through GPI linkage is required for localization and that Dpr–DIP intracellular signaling likely requires transmembrane co-receptors.

## INTRODUCTION

Cell Surface Proteins (CSPs) allow a cell to interact with neighboring cells and the extracellular matrix and interpret chemical cues from its environment. CSPs can be attached to cells through hydrophobic transmembrane domains that traverse the lipid bilayer or via post-translational modifications that anchor the protein in the plasma membrane. One such modification is the addition of a glycosylphosphatidylinositol (GPI) anchor, which is covalently linked to the C-terminus of the protein and embeds its hydrophobic acyl chains into the outer leaflet of the cell membrane.^1–3^ GPI anchors are attached at the ω site, typically a small amino acid, flanked upstream by an unstructured region and downstream by a stretch of hydrophobic residues.^4^ The GPI anchor is added in the ER lumen during protein synthesis, and the mature protein is trafficked to the cell membrane through the secretory pathway.^5^ GPI-anchored proteins cannot signal into the cells directly, as the anchors do not physically reach into the cell interior. Thus, the interaction of GPI linked proteins with intracellular signaling pathways likely requires engagement of transmembrane co-receptors able to transduce the signal inside the cell.

The human genome is predicted to encode at least 129 GPI-anchored proteins.^6^ These include molecules implicated in a variety of functions in the nervous system including axon guidance and synaptic adhesion (reviewed in Um and Ko, 2017)^7^. Abrogation of GPI anchoring and defects in biosynthesis of GPI anchors have been implicated in various diseases including central nervous system disorders.^8, 9^

Computational tools to predict GPI-anchored proteins have been unreliable due, in part, to small training sets and similarity of features associated with GPI anchors and transmembrane domains. Compounding issues with predictions, the membrane-anchoring status of a particular protein cannot simply be deduced from its homology with other proteins; protein families may include members that can be transmembrane, GPI anchored and secreted. For example, the Semaphorin family of key axon guidance cues includes members that are secreted (classes 2 and 3), transmembrane (classes 1, 4, 5, 6) and GPI anchored (class 7).^10^ Sema7A is GPI anchored and involved in axonal outgrowth,^11^ synaptic pruning,^12^ and integrin-dependent stimulation of immune cells.^13^ Ephrins, the membrane-tethered ligands for the receptor tyrosine kinases, Ephs, are divided into GPI anchored ephrin-A and single-pass transmembrane ephrin-B classes and act by binding their cognate EphA and EphB receptors, respectively.^14, 15^ Interestingly, GPI-anchored Ephrin-A5 has been shown to transmit an intracellular signal despite the lack of intracellular region; it partitions into caveolae-like microdomains and engages Fyn protein tyrosine kinase signaling upon interaction with an externally applied soluble form of EphA5 receptor.^16^

The IgLONs, a five-member family of conserved mammalian CSPs, are GPI anchored and function in neurite outgrowth and synaptogenesis.^17–20^ IgLON-mediated signaling remains poorly understood although one member, Negr1, is cleaved by an ADAM-family protease and binds to an FGF receptor to stimulate dendritic arbor growth.^21^ The *Drosophila melanogaster* orthologs of the IgLONs, the Dpr–DIP family of CSPs,^22^ have been implicated in a variety of functions, including synapse specificity and partner preference,^23–28^ axonal pathfinding and fasciculation,^29–31^ cell fate determination,^25^ cell survival,^24, 28, 32^, and behavior.^33^ Structurally, the ectodomains of Dprs and DIPs consist of two and three immunoglobulin (Ig) domains, respectively,^24, 34^ similar to IgLONs which have three Ig domains and interact in a structurally similar manner to Dprs and DIPs.^22, 35^ However, how these proteins are linked to the cell membrane has not been determined. Here, we tested whether members of the *Drosophila* Dpr/DIP family are GPI anchored. We observed that all Dprs and DIPs have a GPI anchor; this was not expected as only a few family members are predicted to have GPI anchors. Treatment with GPI-specific phospholipase-C (PLC) causes shedding of Dprs and DIPs from the surface of insect cells and live fly tissue; this cleavage is GPI anchor-dependent, as it is lost when the GPI anchor site is replaced with a transmembrane domain of a CD4 glycoprotein. Additionally, cleavage of GPI anchors also abolishes Dpr–DIP-mediated cell aggregation. Finally, replacing the GPI anchor with a transmembrane domain perturbs presynaptic and postsynaptic localization of these CSPs. Together these findings suggest that GPI anchors of Dpr/DIP contribute to their localization and function.

## RESULTS

### Some Dprs and DIPs are predicted to be GPI anchored

The Dprs and DIPs bind selectively with one another via their ectodomains, forming an elaborate network of homo- and heterophilic interactions (Figure 1A). Their multifunctional roles in neural circuit development partially relies on their abilities to act as cell adhesion molecules. However, whether and how Dprs and DIPs signal intracellularly has not been examined. Determining if they are transmembrane (TM) or GPI anchored proteins would shed light on their potential signaling mechanisms.

**Figure 1.**
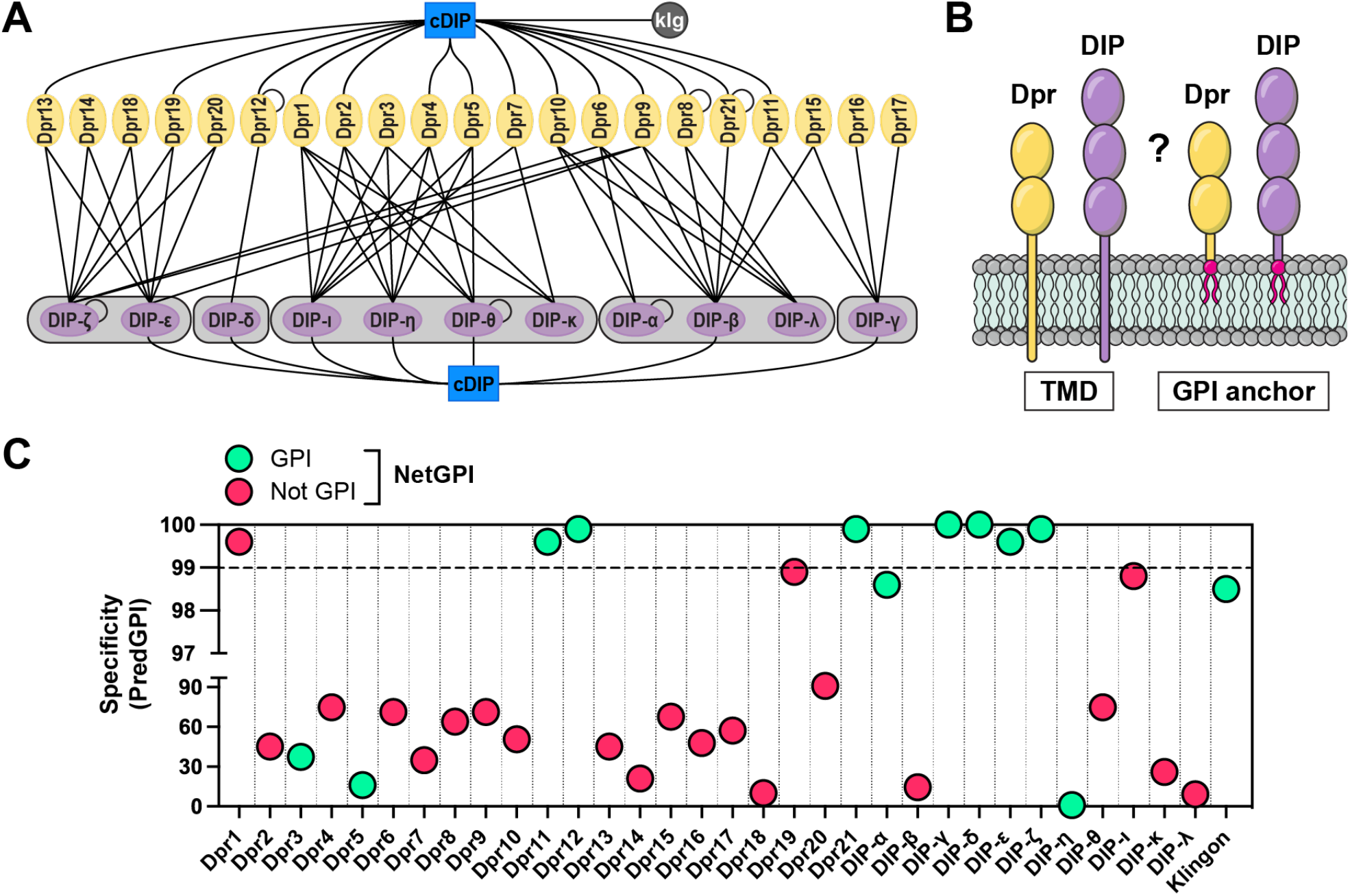
GPI anchoring predictions for the Dpr and DIP families of CSPs. (A) Dpr-DIP interactome. Lines indicate interactions determined in previously published *in vitro* studies. cDIP, common DIP-interacting protein; Klg, Klingon. (B) Two potential membrane-anchoring modes of Dprs and DIPs. TMD, transmembrane domain; GPI, glycosylphosphatidylinositol. (C) GPI anchor predictions using PredGPI and NetGPI. Specificity values as calculated by PredGPI were plotted for each protein. Proteins with specificity values above 99% are considered GPI-anchored. For NetGPI, positive predictions are shown as green and negative are shown as red.

To gain insight into how Dprs and DIPs associate with the cell membrane we used two transmembrane region prediction tools, Phobius^36^ and DeepTMHMM^37^. We first removed N-terminal signal peptides recognized by SignalP-6.0^38^ to avoid their classification as TM domains, which is an inherent challenge for TM region prediction tools due to the hydrophobic nature of both types of sequences. Phobius predicted 15 of the Dprs and DIPs and Klingon to have C-terminal TM regions while DeepTMHMM indicated only Dpr9 as a TM protein (Figure S1).

Next, we hypothesized that those Dprs and DIPs that were not classified as transmembrane may be anchored to the membrane via GPI modification (Figure 1B). We used two GPI signal prediction tools, PredGPI^39^ and NetGPI^40^. PredGPI predicts the likelihood that a protein is GPI anchored and the position of the ω site—the residue to which the lipid anchor is attached. PredGPI is based on Hidden Markov Model, trained on experimentally validated GPI anchored proteins, and produces minimal false positives while correctly identifying 89% of known GPI-containing proteins. NetGPI is a more recently developed prediction software that uses neural networks to predict GPI anchoring, the ω site, and the likelihood of the selected position being correct. NetGPI was reported to slightly outperform PredGPI.^40^

We examined all 32 members of the Dpr/DIP family and Klingon, an immunoglobulin superfamily (IgSF) protein previously demonstrated to be GPI anchored,^41^ with PredGPI and NetGPI and the majority of these CSPs were predicted to lack GPI anchors (Figure 1C and Table S2). PredGPI indicated 8 Dprs and DIPs (25%) as likely GPI linked (specificity above 99%) and Klingon as not GPI anchored as it fell under the threshold for positivity with its specificity value of 98.5%. NetGPI predicted 11 Dprs and DIPs (34%) to be GPI anchored and correctly classified Klingon. Among the proteins predicted to have a GPI modification by NetGPI, 67% were also predicted by PredGPI. Surprisingly, Dpr3, Dpr5, and DIP-η were classified by NetGPI as likely GPI anchored, but obtained very low specificity index values of 37.30%, 16.00%, and 1.00%, respectively with PredGPI.

Overall, 12 Dprs and DIPs were predicted to be GPI anchored by either PredGPI or NetGPI (Figure S2). 15 Dprs and DIPs were predicted to have TM helices, and five of these CSPs were also predicted to have a GPI anchor (Dpr5, Dpr11, DIP-γ, DIP-ζ, and DIP-η). Ten Dprs and DIPs were not predicted to be GPI linked nor to contain a TM helix (Dpr4, Dpr6, Dpr10, Dpr13, Dpr15, Dpr16, Dpr17, DIP-θ, DIP-ι). Thus, using prediction tools alone did not unequivocally classify how these CSPs are anchored to the cell.

### Dprs and DIPs are anchored to the membrane via glycophosphatidyl linkages

To complement the prediction tools and examine the membrane anchoring mechanism(s) of all Dprs and DIPs, we setup an S2 cell culture pipeline with V5-tagged, full-length proteins (see Table 1 and the methods section for details). Duplicate S2 cultures were established for each CSP, and one culture was treated with the GPI cleaving enzyme Phospholipase C (PLC). Supernatant and cell fractions were collected and used for western blot analyses to determine if the CSP was cleaved by PLC. The GPI-anchored protein Klingon was used as the positive control and the secreted cDIP and transmembrane proteins Roughest (Rst) and Kirre served as negative controls.^24, 41–43^ If the CSPs are GPI anchored, we expect an increase of protein in the supernatant and a concomitant decrease in the cell fraction after PLC treatment (Figure 2A). Remarkably, all Dprs and DIPs displayed these trends, suggesting that, like their vertebrate orthologs, the IgLONs, Dprs and DIPs are GPI anchored (Figure 2B).

**Figure 2.**
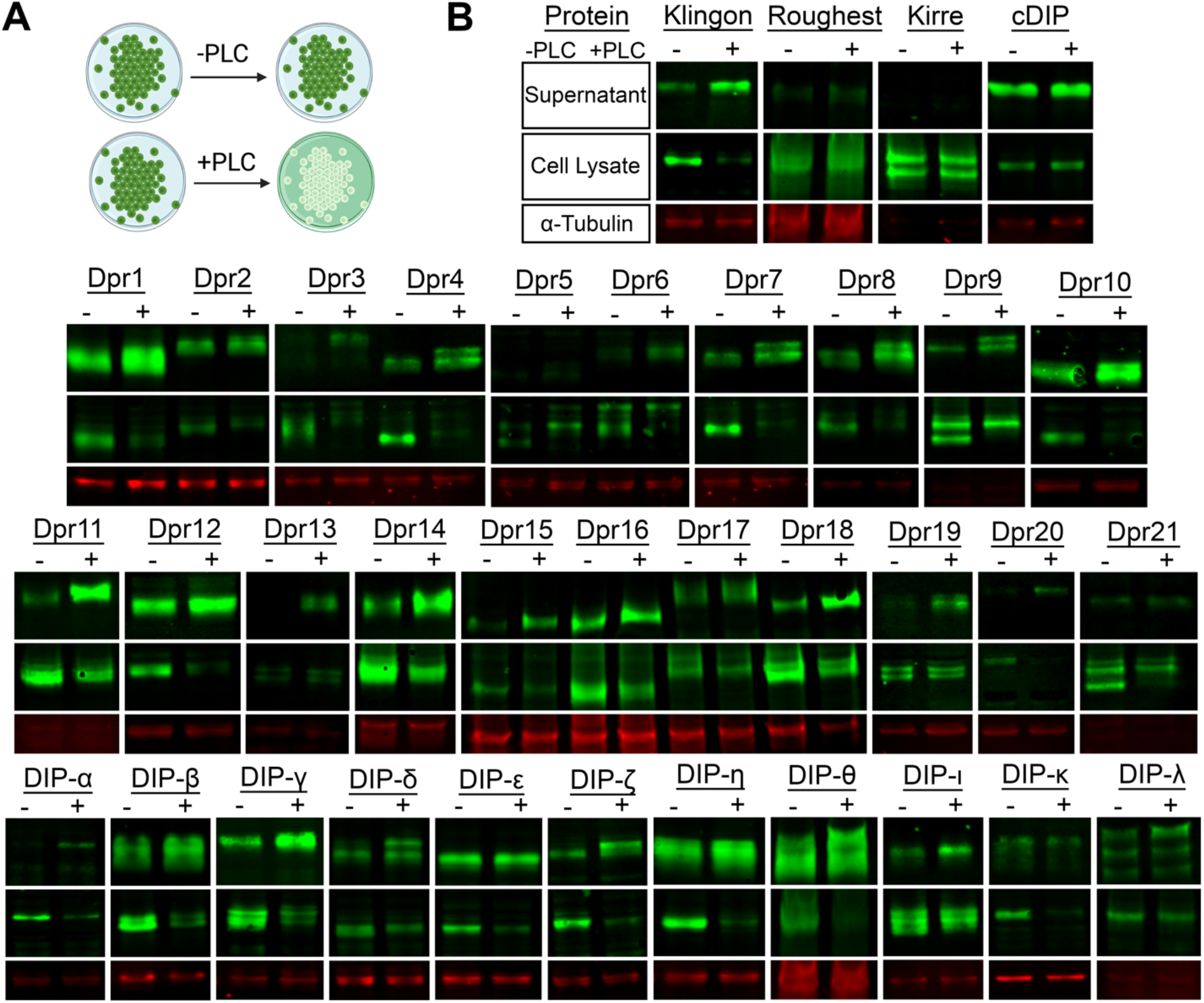
PLC treatment cleaves Dprs and DIPs from S2 cells, as observed by western blotting. (A) Schematic for PLC cleavage assay on S2 cells, when a CSP is GPI anchored. Green color indicates where the GPI-anchored protein is (on cells or in culture media). (B) Western blots of the supernatant fraction, cell lysate, and Tubulin-α loading control for with (+) and without (–) PLC treatment for all members of Dpr/DIP subfamilies. See key in the top row for B.

We next used flow cytometry as an orthogonal method to demonstrate the GPI anchoring of Dprs and DIPs. We expressed the same, N-terminally V5-tagged constructs of three Dprs and three DIPs in Sf9 cells using baculoviral infection. The cultures expressing individual Dprs and DIPs, as well as the negative control, Rst, were split into two samples: one was treated with PLC, and the other served as an untreated control. Both samples were stained with fluorescent antibodies against V5 tag and the relative levels of proteins on the surface of PLC-treated and untreated cells were assessed using a flow cytometer. All DIPs tested, DIP-α, DIP-β, and DIP-γ, showed a significant decrease in protein levels on the surface of Sf9 cells after PLC treatment (Figure 3A-C). Similarly, all Dprs tested, Dpr10, Dpr11, and Dpr21, were mostly cleaved off the cell surface by PLC (Figure 3D-F). In contrast, the level of Rst on the cell surface remained unaffected by PLC cleavage (Figure 3G-H), as expected for a transmembrane protein. The differential extent of cleavage between all tested Dprs and DIPs could be explained by the potentially different accessibility of the GPI cleavage site for each of the proteins. The results obtained using flow cytometry corroborate the observations made using western blot analyses. Together, these findings demonstrate that Dprs and DIPs are GPI anchored.

**Figure 3.**
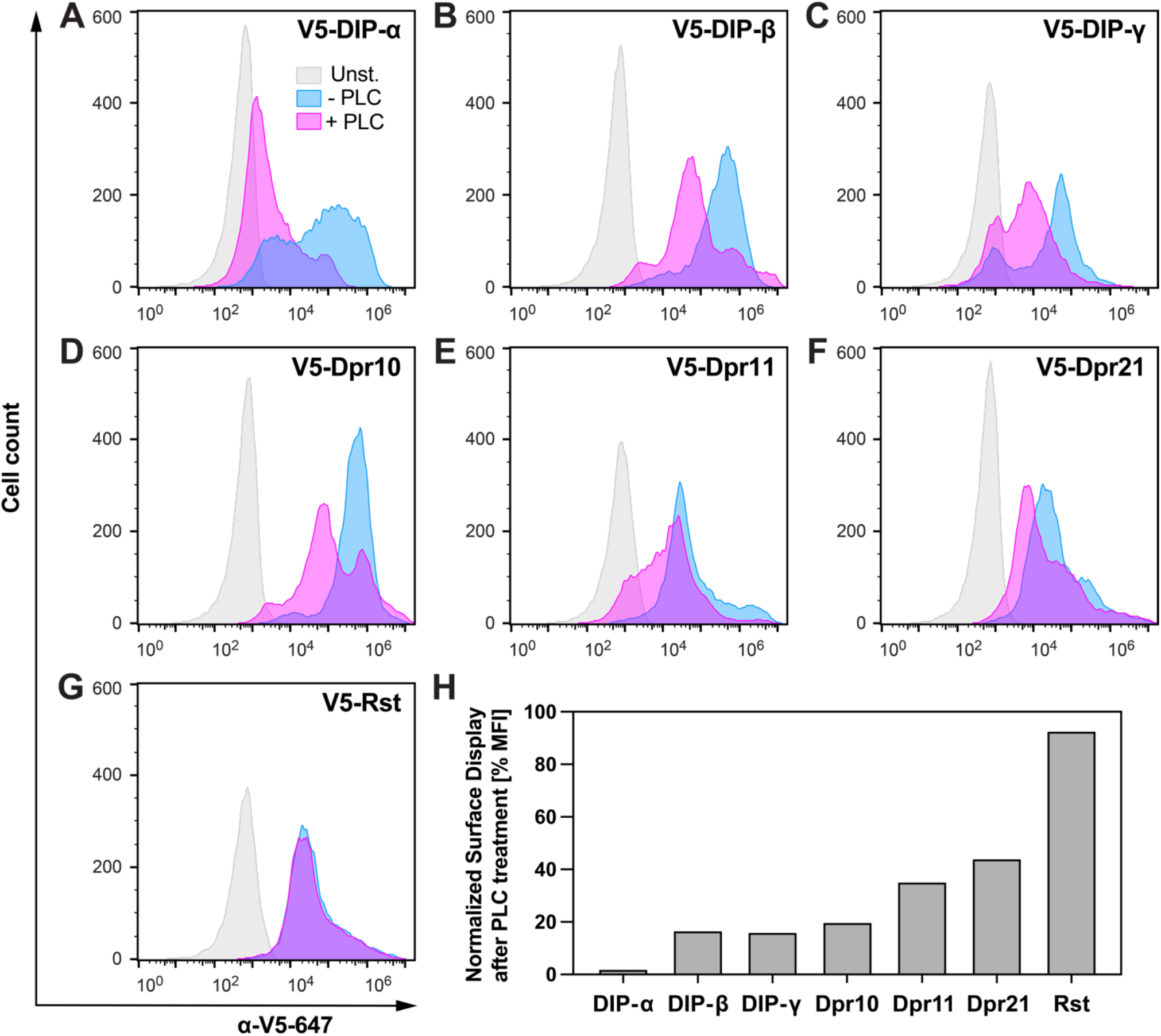
Surface display of Dprs and DIPs is reduced by PLC treatment, as observed by flow cytometry. (A-G) Histograms showing fluorescence levels of baculovirus-infected, unstained Sf9 cells (gray), cells infected and stained with anti-V5-Alexa Fluor 647 antibody (blue), and cells infected, treated with PLC and stained with anti-V5-Alexa Fluor 647 antibody (magenta). (H) Remaining Dpr and DIP levels on Sf9 cell surface following PLC treatment, normalized as the ratio of median fluorescence intensity (MFI) for PLC-treated and not treated cells after subtraction of the background MFI value of unstained cells.

### GPI anchor cleavage eliminates Dpr–DIP-mediated cell aggregation

Dprs and DIPs are thought to mediate cell adhesive interactions to accomplish many of their roles in the nervous system.^32^ A powerful in vitro approach to study cell adhesion interactions is the cell aggregation assay.^44–46^ Here, we combined cell aggregation and PLC assays to test whether cleavage of GPI anchors abrogates cell adhesion mediated by Dprs and DIPs. We used Sf9 cells infected with baculoviruses encoding mScarlet or EGFP followed by V5-tagged, full-length Dpr or DIP, respectively, separated by a P2A peptide. We confirmed the expression of Dprs and DIPs using western blots with an anti-V5 antibody (Figure S3). Cultures expressing individual Dprs with mScarlet were mixed with cultures expressing cognate DIP partners and EGFP. We predicted that, if Dprs and DIPs are GPI-anchored cell adhesion molecules, Dpr– DIP interactions would lead to aggregation between the respective cells, and application of PLC would break up the aggregates. Among the four pairs of cognate Dprs and DIPs tested, DIP-α and Dpr6, DIP-β and Dpr10, and DIP-γ and Dpr11 induced robust aggregation of Sf9 cells (Figure 4A-E). DIP-β and Dpr21-expressing cells aggregated significantly less, which was surprising considering their relatively high affinity compared to other Dpr–DIP pairs,^47^ and that the expression levels of DIP-β and Dpr21 were comparable to other proteins tested here. One reason for the reduced aggregation may be weak homophilic Dpr21 interactions (KD ∼ 50 uM) on the same cell that may prevent efficient heterophilic interactions with DIP-β between cells, as homophilic and heterophilic complexes use the same interfaces and cannot co-exist.^47, 48^ Nonetheless, the addition of PLC significantly reduced cell aggregation for all four pairs of Dprs and DIPs. These results demonstrate that Dpr–DIP interactions instruct cell adhesion and suggest that Dprs and DIPs are anchored to the cell surface membrane via GPI modifications.

**Figure 4.**
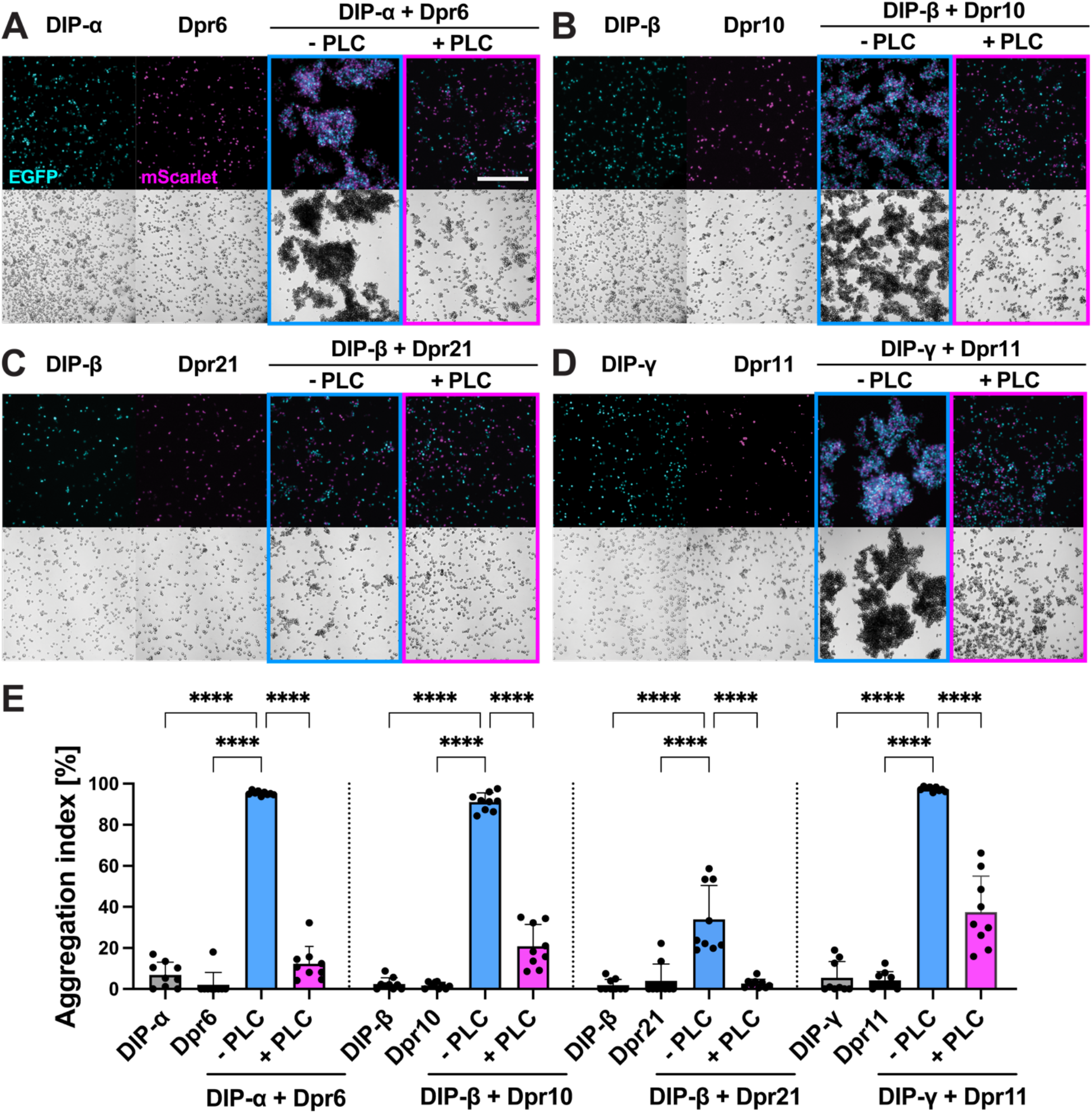
PLC cleavage eliminates Dpr–DIP-mediated cell aggregation. (A-D) Cell aggregation experiments with Sf9 cells. Controls included cultures expressing either DIP or Dpr (first two panels from the left) and cultures expressing DIP or Dpr mixed together but not treated with PLC (-PLC). Cell aggregation was abolished when PLC was added to the mixed cultures 30 minutes after aggregation (+PLC). Scale bar = 500 μm. (E) Cell aggregation index for the samples shown in A-D; ****p<0.0001; one-way ANOVA.

### Dpr and DIP proteins are GPI anchored in vivo

Having demonstrated that Dprs and DIPs are GPI anchored in vitro, we wanted to determine if the same modification occurs in fly tissue. We expressed tagged forms of Dprs and DIPs in the neuromuscular circuit using the GAL4-UAS system. The larval *Drosophila* neuromuscular junction (NMJ) is an excellent system to examine cell surface proteins, as the circuit is well characterized and easily imaged. Moreover, Dprs and DIPs are endogenously expressed at the larval NMJ,^49^ and several Dprs and DIPs are implicated in NMJ development. DIP-α and Dpr10 are known to function in motor axon pathfinding and innervation,^23, 31^ and DIP-γ and Dpr11 regulate synaptic growth through the BMP pathway.^24^

To achieve high protein levels in a cell type that would allow for easy visualization of Dprs and DIPs on the cell surface, we expressed proteins in muscles using *Mef2-GAL4* and omitted detergents in the initial steps of our staining protocol. Muscles are highly stereotyped, multinucleated cells that are easily imaged. In third instar larvae, V5-Dpr19 localized to the subsynaptic reticulum (SSR), a network of postsynaptic membrane folds surrounding the innervating presynaptic boutons (Figure 5A-B). After incubation with PLC, nearly all surface V5 signal was lost, indicating that Dpr19 is GPI anchored in vivo (Figure 5M). Similarly, V5-Dpr10 was localized to the SSR and PLC treatment significantly decreased the V5 signal (Figure 5C-D,N). However, unlike Dpr19, significant Dpr10 remained on the muscle surface in puncta surrounding boutons (Figure 5D,D’), suggesting that a population of Dpr10 was inaccessible to PLC or prevented from diffusing away by interacting with other CSPs at these sites. To confirm that loss of the cell surface signal was due to cleavage of the GPI anchor, we generated chimeric transmembrane tethered Dpr10 proteins by replacing the second Ig domain and C-terminus of Dpr10 with the fourth Ig and transmembrane domains of CD4 (a transmembrane protein not endogenously found in flies). As expected, the cell surface abundance of this chimeric Dpr10-CD4 was unaffected after PLC treatment, suggesting that Dpr10 is anchored at the NMJ via GPI modification (Figure 5E-F’,O). Conversely, when we replaced Ig1 of Dpr10 with the first Ig domain of CD4 but retained the second Ig and C-terminus, this chimeric CD4-Dpr10 was efficiently released from the muscle surface by PLC (Figure 5G-H’,P), indicating that some PLC-released Dpr10 was likely retained on the cell surface through its adhesion domain, Ig1, via interactions with endogenous Dprs and DIPs. These results demonstrate that Dpr10 and Dpr19 are GPI anchored in their endogenous tissue.

**Figure 5.**
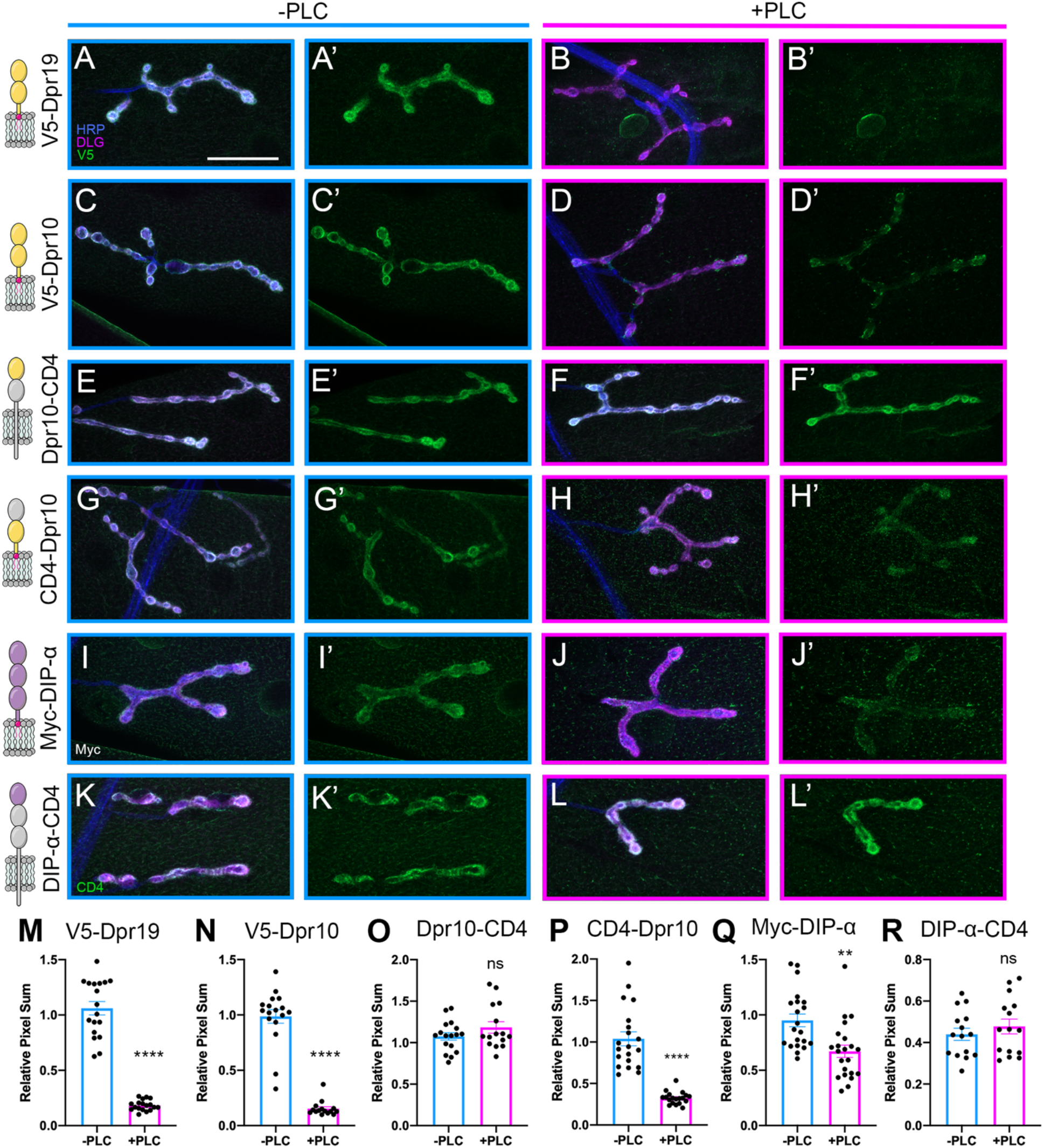
Dpr and DIP proteins are GPI anchored in vivo. (A-L’) Tagged Dprs and DIPs were expressed in muscles, and dissected larvae were treated with PLC and compared to non-treated controls. Cartoon presentations of tagged Dprs and DIPs and their domains are depicted in left column. Surface localized proteins are shown in green, neuronal tissue in blue (HRP), and postsynaptic membrane in magenta (DLG). (A-B) PLC efficiently reduced surface labeling of V5 on muscles expressing V5-Dpr19. (C-D) PLC efficiently reduced surface labeling of V5 on muscles expressing V5-Dpr19. (E-F) PLC failed to reduce labeling of V5 in muscles expressing Dpr10-CD4 chimeras. (G-H) PLC efficiently reduced surface labeling of V5 in muscles expressing CD4-Dpr10 chimeras. (I-J) PLC reduced surface labeling of Myc on muscles expressing Myc- DIP-α. (K-L) PLC failed to reduce muscle labeling of CD4 on muscles expressing DIP-α-CD4 chimeras (M-R) Quantification of experiments shown in A-L’. Scale bar = 50 μm.

Next, we examined two DIPs, DIP-α and DIP-ζ. Although neither is endogenously expressed in muscles,^49^ an ectopically expressed DIP-α variant localized to the SSR (Figure 5I). Like Dpr10 and Dpr19, DIP-α was released from the muscle surface after PLC treatment (Figure 5I-J’,Q). However, cleavage was less efficient, and DIP-α puncta formed on the muscle surface and around boutons after treatment. Similar to Dpr10, replacing Ig2-Ig3 and the C-terminus of DIP-α with CD4 blocked the PLC-mediated release from the muscle surface (Figure 5K-L’,R). Strikingly, DIP-ζ did not localize to the SSR, and PLC treatment resulted in increased DIP-ζ puncta across the muscle surface (Figure S4A-C). Overall, these data show that PLC treatment affects Dpr10, DIP-α and - ζ attachment and localization in vivo and support our model that Dprs and DIPs are GPI anchored CSPs.

### GPI anchor alters localization in neurons and muscle cells

GPI modifications have been shown to contribute to the subcellular localization of CSPs.^50, 51^ Thus, we examined localization of ectopically expressed wild type and chimeric Dpr10 and DIP-α in their endogenous tissues. DIP-α is expressed in a subset of motor neurons called the Is type and localizes to the axon terminals.^23, 24, 49^ Using a Is motor neuron-specific driver, *A8-GAL4*,^27, 52^ we confirmed that Myc-DIP-α localizes to puncta in the motor axon terminal (Figure 6A-B’). However, when the GPI anchor was replaced with a TM domain, DIP-α was redistributed and fewer puncta formed, suggesting that clustered presynaptic localization is partially due to the GPI anchor (Figure 6C-D’). These changes were confirmed by quantifying DIP-α puncta in the terminal three boutons in each condition: In order to determine the nature of this localization difference, samples were thresholded and collapsed to a binary representation (Figure 6B,D), and then the number of particles in and around the three terminal boutons of each arbor were quantified. Myc-DIP-α formed more and smaller puncta compared to DIP-α-CD4 (Figure 6I-J), suggesting that the GPI anchor contributes to the subcellular localization of DIP-α in vivo.

**Figure 6.**
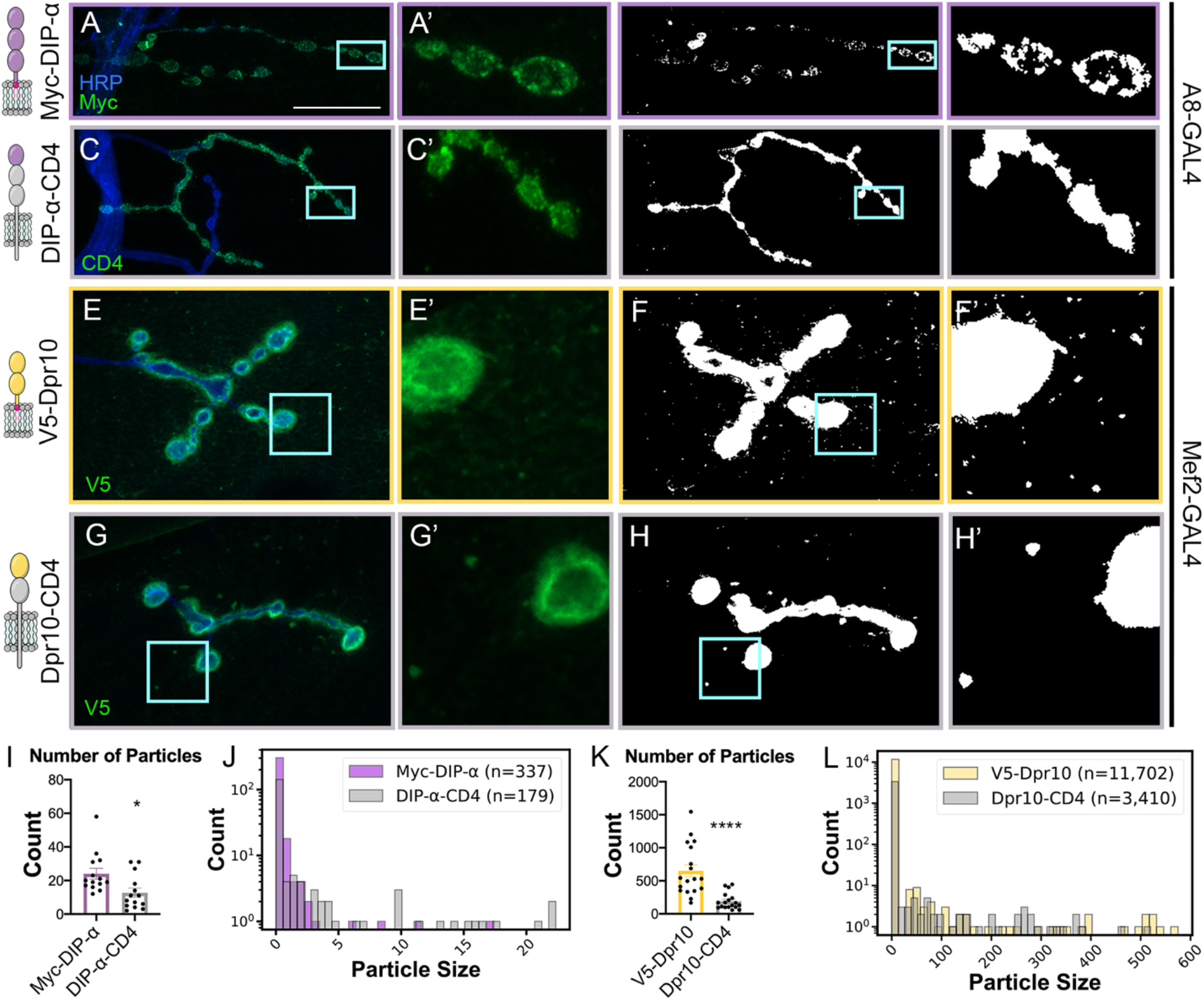
GPI anchor alters localization of DIP-α in neurons and Dpr10 in muscles. (A-H’) Cartoon depictions of constructs are shown in left column. Surface localized proteins are shown in green, neuronal tissue in blue (HRP). Scale bar = 50 μm. (A-B) Tagged DIP-α and DIP-α-CD4 were expressed in Is motor neurons. (A-A’) *A8-GAL4* driving Myc-DIP-α leads to punctate surface localization outside the boutons. (B-B’) Binary thresholded image of A. (C-C’) *A8-GAL4* driving DIP-α-CD4 leads to localization of DIP-α throughout the Is motor neuron terminal. (D-D’) Binary thresholded image of C. (E-E’) *Mef2-GAL4* driving V5-Dpr10 leads to punctate surface localization. (F-F’) Binary thresholded image of E. (G-G’) *Mef2-GAL4* driving Dpr10-CD4 leads to loss of Dpr10 throughout the muscle surface. (H-H’) Binary thresholded image of G. (I,K) Particles counted for experiments depicted in A-D and E-H, respectively. (J, L) Histogram showing counts for particle size distribution.

As described above, expressing Dpr10 in muscles results in Dpr10 localization at the SSR, the membrane folds that form around in presynaptic boutons. However, additional puncta are observable away from boutons on the muscle surface (Figure6 E-H’) and these puncta were significantly reduced when compared to the localization of the chimeric DIP-α containing a transmembrane helix (Figure6 K-L). Taken together, GPI anchors of DIP-α and Dpr10 contribute to both their pre- and postsynaptic localization, respectively.

## DISCUSSION

Dprs and DIPs have been implicated in many aspects of neural circuit development. However, we lack a systematic analysis of how these CSPs are anchored to the cell membrane to provide insights into their signaling mechanisms. Here, we utilize several in vitro and in vivo approaches to demonstrate that all Dprs and DIPs are attached to the cell membrane through GPI anchors. In tissue, we show that GPI anchors contribute to subcellular clustering on the presynaptic neuronal membrane. Our findings, that all Dpr and DIPs are GPI anchored, suggest that Dpr/DIP-mediated functions must be achieved either by merely tethering two cells together and/or by signaling through a co-receptor.

### Predicting GPI anchors

Predicting GPI anchors is intrinsically challenging for multiple reasons. First, the features that predispose the protein to GPI anchor attachment are not very well defined. Second, the ω site is flanked downstream by a stretch of hydrophobic amino acids that can resemble a single-pass transmembrane domain. Third, it is estimated that the animal genomes may each encode over 100 GPI anchored proteins, but only a fraction of those have been experimentally validated, resulting in limited training datasets for prediction tools. Consequently, both currently annotated GPI-anchored proteins and the predictions of such proteins likely represent an underestimate of the actual number. Indeed, a high number of false negatives was observed in this study, as only one-third of Dprs and DIPs were predicted to be GPI anchored by the latest available algorithms in PredGPI and NetGPI. Conversely, the TM prediction tool Phobius predicted 15 out of 32 Dprs and DIPs and Klingon as TM proteins, suggesting this software may not be robust at discriminating between the two membrane targeting motifs. This is not surprising, however, considering the similar chemical nature of TM helices and GPI signals. The newest version of TMHMM, DeepTMHMM, a deep learning-based algorithm, performed significantly better than Phobius, classifying only Dpr9 as a TM protein. Unexpectedly, one-third of Dprs and DIPs were neither predicted to be GPI anchored nor to have a TM helix. Together our findings demonstrate that currently available GPI signal prediction tools can provide useful preliminary insights, especially in combination with TM prediction software. However, experimental validation should be the primary way of assessing GPI anchor presence, when possible, until better prediction tools are developed.

### Dpr and DIP clustering is aided by their GPI anchors

CSP localization is often critical for their function. Here, we demonstrate that DIP-α, Dpr10, and Dpr19 localized to the postsynaptic membrane when expressed in muscles. Moreover, punctate Dpr10 localization on the muscle surface was diminished when we replaced the GPI anchor with a TM helix suggesting that this modification contributes to its localization. Similar results were observed for punctate DIP-α in presynaptic Is arbors. GPI-anchored proteins have been reported to localize to lipid rafts—domains in the lipid bilayer that have increased levels of sterols and are therefore slightly thicker and less mobile than the surrounding membrane.^53^ Although still somewhat controversial, these domains are thought to scaffold proteins by increasing the local concentration of proteins that preferentially localize to lipid rafts such as GPI-anchored proteins, possibly including Dprs and DIPs within these rafts.

The PLC cleavage of Dpr10 and DIP-α from muscles was incomplete, leaving residual punctate signal around the boutons and on the muscle surface unlike Dpr19 that was almost completely lost. These differences in PLC cleavage efficiencies may be due to different local interactions of Dprs and DIPs with various proteins or lipids in the membrane. Alternatively, we provide evidence suggesting that cleavage is complete and the observed signal is due to the cleaved CSPs binding nearby proteins and preventing release from the tissue.

### Dpr–DIP-mediated cell adhesion depends on their GPI anchors

*Trans*- and *cis*-interactions between Dprs and DIPs drive their roles in the nervous system. Dprs and DIPs may act similar to other cell adhesion molecules (e.g., integrins or cadherins) by clustering to generate avidity and strengthen cell-cell contacts during axonal fasciculation or synapse formation. We examined four Dpr–DIP pairs and all showed robust aggregation that was susceptible to PLC treatment, suggesting that the GPI anchors in Dprs and DIPs are accessible by PLC at the cell-cell contact sites. This phenomenon could have profound consequences in vivo for modulating cell adhesion, provided that specific lipases and proteases are present under physiological conditions at sufficient concentrations and exhibit high enough substrate turnover to cause structural changes to cell-cell contacts.

Shedding of cell surface molecules by hydrolytic enzymes like proteases and lipases has been implicated in various signaling mechanisms. Examples include activation of fibroblast growth factor 2 (FGF-2) via the release of soluble syndecan-1,^54^ potentiation of β-catenin signaling upon cleavage of neuronal cadherin (N-cadherin),^55^ modulation of Nodal signaling via GPI cleavage of CRIPTO,^56^ and bi-directional regulation of the Notch pathway through proteolysis of Notch or its ligands.^57–60^ GPI-anchored proteins are unable to directly signal intracellularly so acting as soluble factors for other receptors, either in *cis* or *trans*, could be a conceivable mode of action. A subset of Dprs^61^ and DIPs were identified in a secretome screen as soluble factors in the hemolymph of larvae (Personal Communication, Norbert Perrimon and Justin Bosch), but the roles of these soluble variants are unknown. In addition to potential functions of soluble forms of these CSPs in the extracellular space, it would be interesting to explore if their shedding in vivo could affect Dpr–DIP-mediated cell adhesion, and have consequences for the structure and function of synapses or other specialized sites of cell-cell contact, as shown for some CAMs.^55, 62–66^

### GPI anchors in neural circuit assembly

GPI-anchored proteins are critical for many physiological processes, including neural circuit formation. These proteins can be cleaved by lipases or proteases as part of their signaling mechanism. For example, the GPI-anchored protein RECK regulates motor neuron differentiation in mammals by inhibiting Notch signaling; only when RECK is cleaved and released from the membrane can the ADAM10 metalloprotease access and cleave the Notch ligand thereby promoting differentiation.^67^ In addition, IgLONs, the vertebrate orthologs of Dprs and DIPs, are shed from neurons via ADAM10 to promote neuronal growth of nearby neurons.^19, 21^ The presence of GPI anchors in Dprs and DIPs suggests that their biology and signaling may be conserved with IgLONs. Finally, if the GPI-anchored protein is not cleaved, it can still signal through a co-receptor that traverses the cell membrane. To our knowledge, no Dpr or DIP co-receptors have been published and this provides an intriguing future avenue of research.

## Acknowledgments

We would like to thank Szymon Kordon, Ruiling Zhang, Gabriel Carmona Rosas, Indya Weathers, Stephanie Dunning, Veera Anand, and Yupu Wang as well as members of the Carrillo and Özkan Labs for support and feedback. This work is supported by NINDS R01 NS123439 and a UChicago Faculty Diversity Grant to R.A.C., NINDS R01 NS097161 to E.Ö., and NIGMS T32 GM007183 and NINDS FS31 NS120458 to M.L-R.

## METHODS

### Sequence selection and molecular cloning

cDNAs of most of the full-length Dprs and DIPs used in the study were obtained from Drosophila Genomics Research Center (DGRC). The remaining full-length sequences were generated through the extension of existing ectodomain constructs subcloned by our groups.^34^ Among the DGRC clones, the sequence of Dpr15 required modification to remove the insertion within the Ig2 domain to preserve its structural integrity. The source, isoform, and identification number of each used sequence can be found in Table S1. Sequences of full-length Dprs and DIPs with an N-terminal V5 tag were cloned into the pMT/BiP/V5-His A vector (Invitrogen) for expression in S2 *Drosophila* cells for PLC cleavage assay or into the pAcGP67 baculovirus transfer vector (BD Biosciences) for expression in Sf9 cells for cell flow cytometry and cell aggregation experiments. For cell aggregation, the constructs additionally included EGFP or mScarlet separated from V5-tagged Dprs or DIPs by the P2A sequence.

### Predictions of GPI anchors and transmembrane helices

Predictions of GPI-linkage were carried out using PredGPI^39^ and NetGPI^40^ tools available online. For predictions using PredGPI the general model was used. Predictions of transmembrane regions were performed using Phobius^36^ and DeepTMHMM^37^. Before analysis in Phobius and DeepTMHMM, the sequences of signal peptides predicted with SignalP-6.0^38^ were removed.

### *Drosophila* reagents, dissections, and immunohistochemistry

All flies were kept at 25°C except when otherwise noted. Crosses were set at medium density and crawling third instar larvae were dissected as before^49^ in Phosphate Buffered Saline (PBS, pH 7.4) on Sylgard dishes. Briefly, after fixing for 30 minutes in 4% paraformaldehyde, fillets were washed 3 times for 10 minutes in PBST (PBS + 0.05% Triton X-100, pH 7.4). Larval fillets were then blocked in 5% Normal Goat Serum in PBST. Samples were incubated in primary antibodies diluted in block and overnight at 4°C, extensively washed in PBST, and incubated with secondary antibodies for 2 hours at room temperature before the final washes in PBST. All washes and incubations occurred on nutators. Samples were then mounted in Vectashield (Vector Laboratories) and representative images were collected. HRP was used as a marker for neuronal membranes and DLG was used as a marker for postsynaptic membranes.

The antibodies used for this study are:

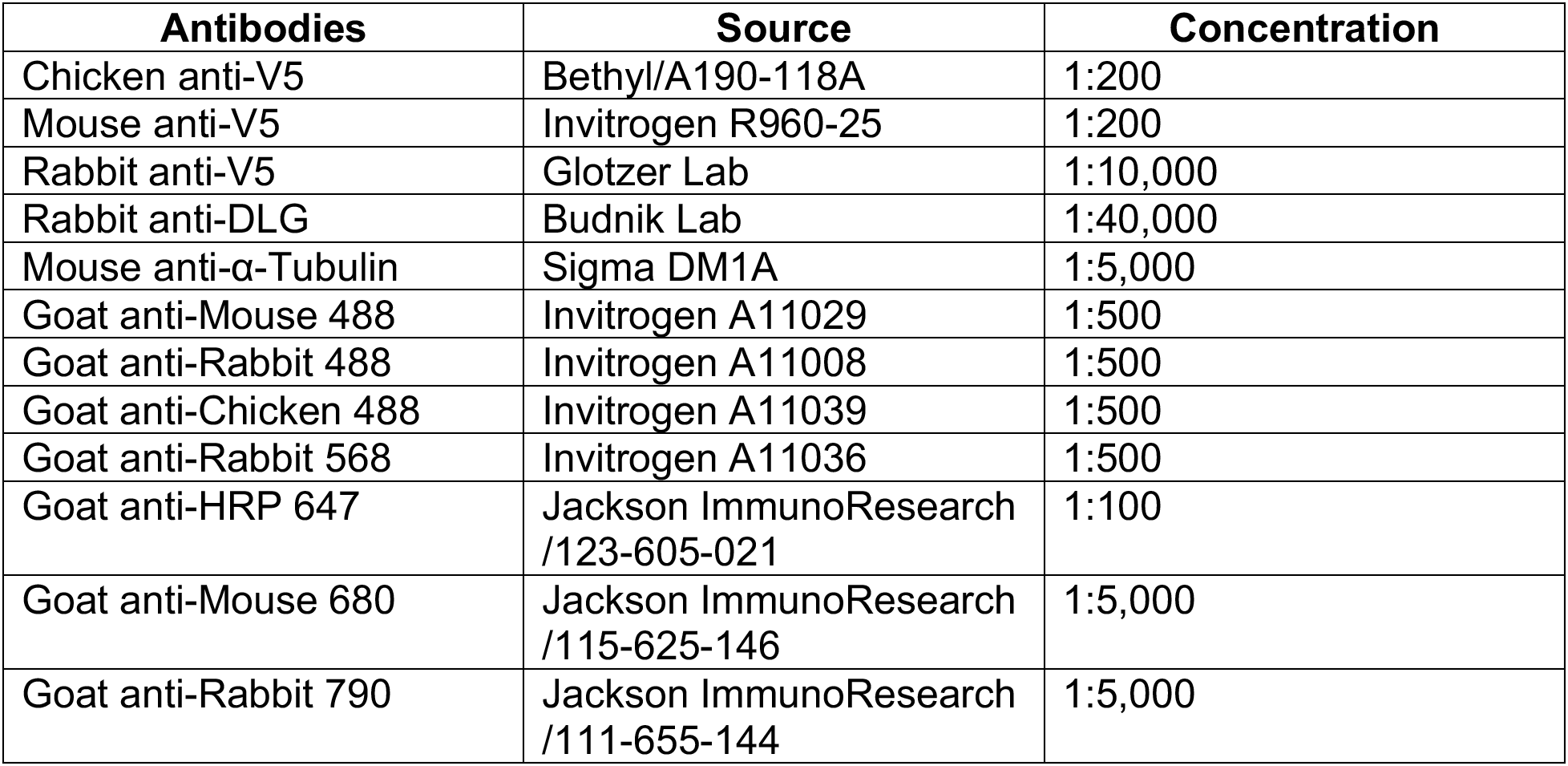

The fly lines used for this study are:

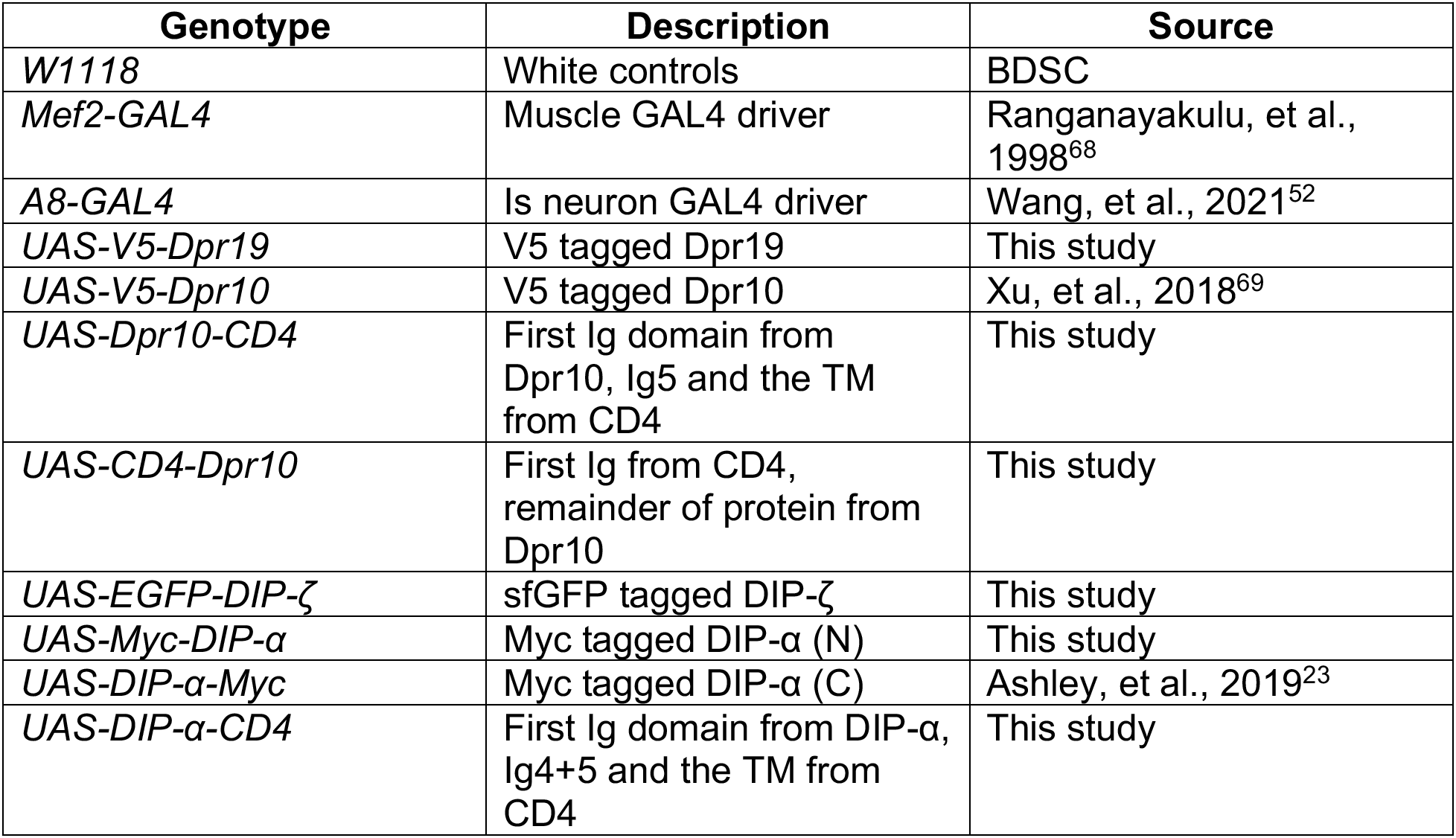

| Genotype | Description | Source |
| --- | --- | --- |
| *W1118* | White controls | BDSC |
| *Mef2-GAL4* | Muscle GAL4 driver | Ranganayakulu, et al., 1998<sup>68</sup> |
| *A8-GAL4* | Is neuron GAL4 driver | Wang, et al., 2021<sup>52</sup> |
| *UAS-V5-Dpr19* | V5 tagged Dpr19 | This study |
| *UAS-V5-Dpr10* | V5 tagged Dpr10 | Xu, et al., 2018<sup>69</sup> |
| *UAS-Dpr10-CD4* | First Ig domain from Dpr10, Ig5 and the TM from CD4 | This study |
| *UAS-CD4-Dpr10* | First Ig from CD4, remainder of protein from Dpr10 | This study |
| *UAS-EGFP-DIP-ζ* | sfGFP tagged DIP-ζ | This study |
| *UAS-Myc-DIP-α* | Myc tagged DIP-α (N) | This study |
| *UAS-DIP-α-Myc* | Myc tagged DIP-α (C) | Ashley, et al., 2019<sup>23</sup> |
| *UAS-DIP-α-CD4* | First Ig domain from DIP-α, Ig4+5 and the TM from CD4 | This study |

### PLC cleavage assay with S2 cells

S2 cells were obtained from the Drosophila Genomics Research Center and maintained in Schneider’s Medium (Sigma S0146), supplemented with Insect Media Supplement (Sigma I7267), and 90 U/ml Penicillin plus 90 μg/ml Streptomycin. Cells were maintained at room temperature. For experiments, confluent cells were split 1:2 into 6-well plates and transfected the following day. Complete Schneider’s Media was mixed 2:1 with 250 µg/ml dimethyldioctadecyl-ammonium bromide (DDAB), allowed to mix, and then 500 μl DNA was added for each well to be transfected. Approximately 24 hours after transfection, protein expression was induced by adding 1 mM copper sulfate (CuSO4). Three days later, 2 μl of 100 U/ml phosphatidylinositol-specific phospholipase C (PI-PLC, Life Technologies P-6466) was added to the treatment well, and cells were incubated for 4 hours.

To harvest cells, 2 ml from each well was collected and spun down at 500 x g for 5 minutes. The supernatant was collected and mixed with 6x Loading Buffer (375 mM Tris-Cl, pH 6.8, 9% SDS, 50% glycerol, 0.03% bromophenol blue, 9% β-mercaptoethanol) and boiled for 10 minutes. The cell pellet was washed in PBS and spun down as before. Next, cells were lysed used using a buffer adapted from Bumgarner et al., 2005,^70^ consisting of 50 mM Tris-HCl pH 8.0, 150 mM NaCl, 1% Triton X-100, 5 mM EDTA, and protease inhibitors (one tablet per 50 ml; Pierce A32955). Tubes were incubated on a nutator at 4°C for 30 minutes and then at 37°C for 30 minutes. Tubes were centrifuged at 17,000 x g at room temperature for 15 minutes and mixed with 6x Loading Buffer.

Samples were then run on 12% SDS-polyacrylamide gels using the TGX FastCast system (BioRad 1610175). The samples were transferred overnight onto a nitrocellulose membrane (BioRad 1620115) at a constant current of 40 mA and blocked for one hour in 1% (w/v) Casein Block. Staining with primary and secondary antibodies was performed for 2 hours at room temperature with slight agitation and included washes in PBST after each incubation. Westerns were imaged on a LI-COR Odyssey imager.

### Flow cytometry

Sf9 cell cultures at the density of 2×10^6^ cells/ml were placed in 6-well plates, 3 ml per well. The cells were infected with baculoviruses encoding full-length Dprs and DIPs with an N-terminal V5 tag. Infected cultures were incubated for 48 hours on an orbital shaker at 125 rpm (Thermo MaxQ 430), at 28°C. 0.6 ml of each culture was transferred to a 24-well plate. 4 μl of PI-PLC per well was used for GPI cleavage. 4 µl of PI-PLC storage buffer (20 mM Tris-HCl, pH 7.5, 1 mM EDTA, 0.01% sodium azide, 50% glycerol) was added to control samples. Treatment was carried out for 3 hours on an orbital shaker at 125 rpm, at 28°C. Cultures were spun down for 1 minute at 500 x g. The cell pellets were resuspended in 300 µl of cold PBS, pH 7.4, with 1% BSA. 100 µl of cell suspensions were transferred to a U-shaped 96-well plate. 4 µl of anti-V5-AF647 antibody (R&D Systems, FAB8926R) was used for staining cells for 20 minutes on a shaker at 500 rpm (Thermo Microplate Shaker), at 4°C. Cultures were spun down for 1 minute at 500 x g, at 4°C and washed 3 times with 200 µl of PBS, pH 7.4, with 1% (w/v) BSA. Cells were analyzed using a BD Accuri C6 flow cytometer. 20,000 events were recorded per sample. The results were analyzed using FlowJo Software v10.8.1 (BD Life Sciences).

### Cell aggregation assay with Sf9 cells

Sf9 cultures at the density of 2×10^6^ cells/ml were placed in 6-well plates, 3 ml per well, and infected with baculoviruses encoding (N- to C-terminal) mScarlet, P2A sequence (ATNFSLLKQAGDVEENPGP) and V5-tagged Dpr, or EGFP, P2A sequence, and V5-tagged DIP. This construct design was chosen to leave unmodified C-termini for Dprs and DIPs, as that would be important for testing for a C-terminal GPI anchoring signal. Infected cultures were incubated for 48 hours on an orbital shaker at 125 rpm, at 28°C. Expression of V5-tagged Dprs and DIPs was confirmed using western blots with the anti-V5-AF647 antibody. Cultures were diluted 1:15 in Sf9 complete media (Gibco Sf-900 III SFM, with 10% FBS, 2 mM L-glutamine, and 20 µg/ml gentamicin). 200 µl of cultures expressing Dprs were mixed with 200 µl of cultures expressing DIPs in 24-well plates. The control samples included 200 μl of cultures expressing individual Dprs or DIPs mixed with 200 µl of non-infected Sf9 cultures. Samples were prepared in triplicates for each Dpr–DIP pair and each control culture expressing individual Dprs and DIPs. Cultures were left to aggregate for 30 minutes on an orbital shaker at 125 rpm, at 28°C. 4 μl of PI-PLC per well was used for GPI cleavage. 4 μl of PI-PLC storage buffer was added to control samples. Treatment was carried out for 1 hour on an orbital shaker at 125 rpm, at 28°C. Cultures were imaged in 24-well plates at 5x magnification. Three images were collected for every well. Cell aggregation was quantified using the cell aggregation index, defined as the percentage of the total area occupied by cells that is comprised of aggregates.^71^ The area occupied by cells and aggregates was determined using the ‘Analyze particles’ function in Fiji. Aggregates were defined as particles with areas of at least 2900 µm^2^.

### Tissue PLC experiment

Larvae were filleted, rinsed in PBS, and incubated while shaking for one hour in 1 ml of PBS with or without 1 µl of PI-PLC. Larvae were washed with PBS, fixed, and stained as above with a slight alteration: detergent was only used after staining for Dpr/DIP protein tags. After extensive washing in PBS, anti-DLG was diluted in PBST and incubated with larvae for 2 hours at room temperature. The subsequent washing and secondary antibody staining was as described above. The staining procedure for PLC-treated and untreated preparations was carried out in the same tube so that conditions were identical. Three muscle 4 Ib arbors were imaged per animal.

For quantification, images were collected from each experiment using identical imaging parameters. After measuring pixel sum of three terminal boutons per arbor, signal of the Dpr/DIP was normalized to DLG signal to control for any differences in staining conditions between replicates and because DLG should not be affected by PLC treatment. These values are reported as ‘Relative Pixel Sum’ representing Tag/DLG pixel sum.

### Tissue Localization experiment

For localization experiments, crosses were maintained at 18°C in order to dampen protein expression levels. Samples to be compared were pooled and stained as above, omitting detergents in the first round of staining to label only proteins that were present on the cell surface at the time of fixation.

Once images were collected, z-stacks were generated, and the relevant channel was converted to 8-bit format before a threshold was set to convert image to binary representation of the surface protein stain. The same threshold was used for all images. This image was then used to count particles. For presynaptic localization, particles of and surrounding three terminal boutons per arbor were measured. For postsynaptic localization, particles of the entire image were counted. These manipulations were performed using ImageJ FIJI.^72^

### Statistical Analysis

For tissue PLC experiments and tissue localization experiments, two-tailed Student’s t-test was used to determine statistical significance (Prism 8). Cell aggregation experiments were evaluated using one-way ANOVA (Prism 9).

### Tissue and cell imaging protocol

All in vivo images were obtained on a Zeiss LSM800 confocal microscope with a 40X plan-neofluar 1.3NA objective or 63X plan-apo 1.4NA objective. All images of Sf9 cells were obtained using a Leica THUNDER Imager 3D Cell Culture with Leica N Plan 5x/0.12 PH0 objective.

## SUPPLEMENTAL MATERIALS

**Table S1.**
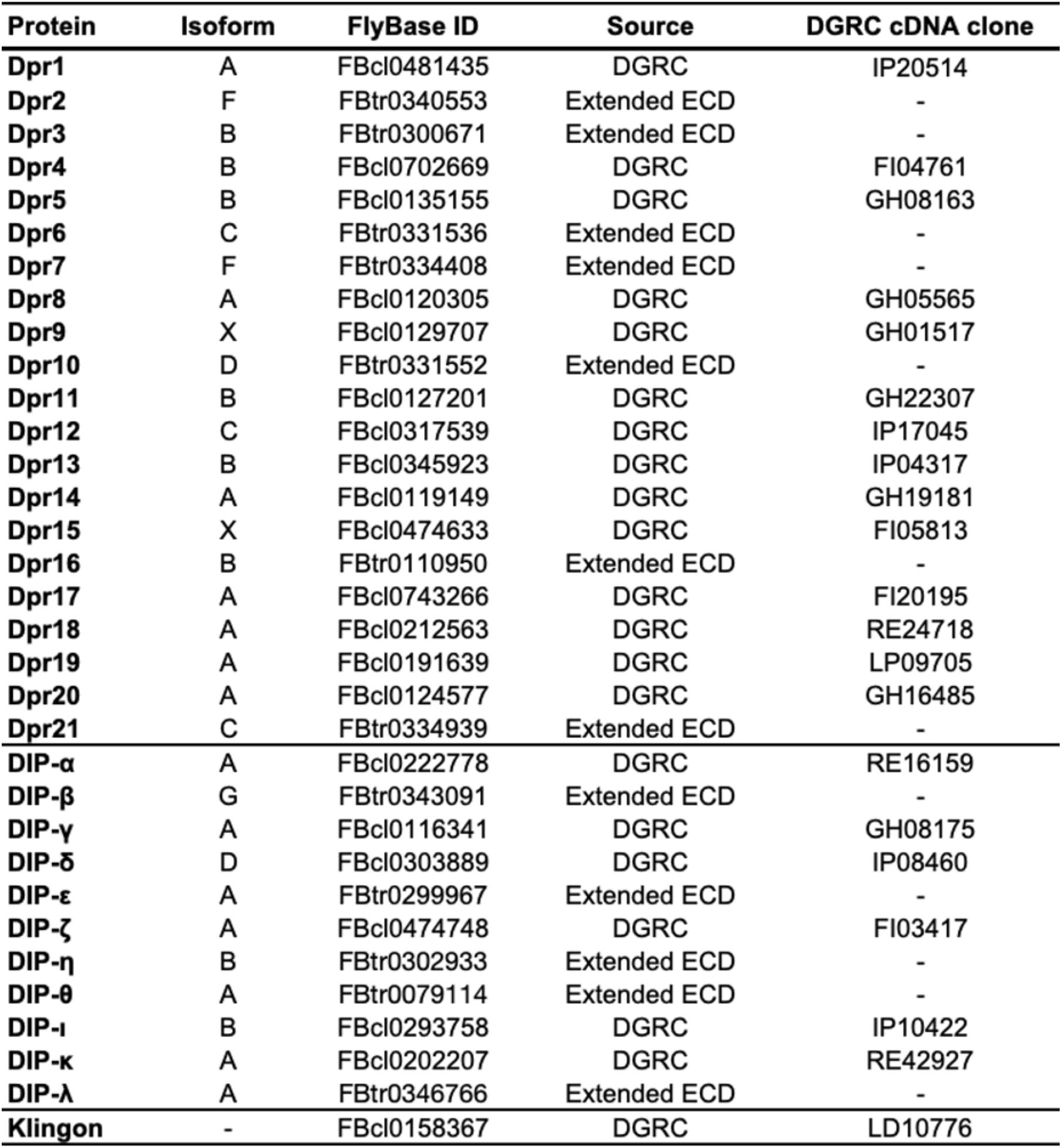
Selection of sequences used in our analyses. X under isoform indicates the exact sequence for the predicted isoform does not exist in Flybase.

**Table S2.** GPI site predictions by PredGPI and NetGPI. Results of GPI-anchor site predictions with PredGPI and NetGPI. Values of specificity index calculated by PredGPI indicate the probability of the presence of a GPI-anchor. Likelihood values for the positive calls from NetGPI pertain to the prediction of the ω site position.

| Protein | PredGPI prediction | Specificity | NetGPI prediction | Likelihood |
| --- | --- | --- | --- | --- |
| Dpr1 | GPI | 99.60% | Not-GPI | 0.975 |
| Dpr2 | Not-GPI | 45.20% | Not-GPI | 0.978 |
| Dpr3 | Not-GPI | 37.30% | GPI | 0.565 |
| Dpr4 | Not-GPI | 74.80% | Not-GPI | 0.813 |
| Dpr5 | Not-GPI | 16.00% | GPI | 0.407 |
| Dpr6 | Not-GPI | 71.20% | Not-GPI | 0.994 |
| Dpr7 | Not-GPI | 34.80% | Not-GPI | 0.508 |
| Dpr8 | Not-GPI | 64.00% | Not-GPI | 0.885 |
| Dpr9 | Not-GPI | 71.20% | Not-GPI | 0.995 |
| Dpr10 | Not-GPI | 50.70% | Not-GPI | 0.989 |
| Dpr11 | GPI | 99.60% | GPI | 0.341 |
| Dpr12 | GPI | 99.90% | GPI | 0.542 |
| Dpr13 | Not-GPI | 45.20% | Not-GPI | 0.922 |
| Dpr14 | Not-GPI | 21.10% | Not-GPI | 0.831 |
| Dpr15 | Not-GPI | 67.70% | Not-GPI | 0.992 |
| Dpr16 | Not-GPI | 47.80% | Not-GPI | 0.305 |
| Dpr17 | Not-GPI | 57.20% | Not-GPI | 0.793 |
| Dpr18 | Not-GPI | 10.10% | Not-GPI | 0.612 |
| Dpr19 | Not-GPI | 98.90% | Not-GPI | 0.352 |
| Dpr20 | Not-GPI | 90.90% | Not-GPI | 0.338 |
| Dpr21 | GPI | 99.90% | GPI | 0.283 |
| DIP-α | Not-GPI | 98.60% | GPI | 0.301 |
| DIP-β | Not-GPI | 14.50% | Not-GPI | 0.941 |
| DIP-γ | GPI | 100.00% | GPI | 0.686 |
| DIP-δ | GPI | 100.00% | GPI | 0.432 |
| DIP-ε | GPI | 99.60% | GPI | 0.45 |
| DIP-ζ | GPI | 99.90% | GPI | 0.643 |
| DIP-η | Not-GPI | 1.00% | GPI | 0.422 |
| DIP-θ | Not-GPI | 74.80% | Not-GPI | 0.606 |
| DIP-ι | Not-GPI | 98.80% | Not-GPI | 0.501 |
| DIP-κ | Not-GPI | 25.90% | Not-GPI | 0.992 |
| DIP-λ | Not-GPI | 9.10% | Not-GPI | 0.802 |
| Klingon | Not-GPI | 98.50% | GPI | 0.486 |

**Figure S1.**
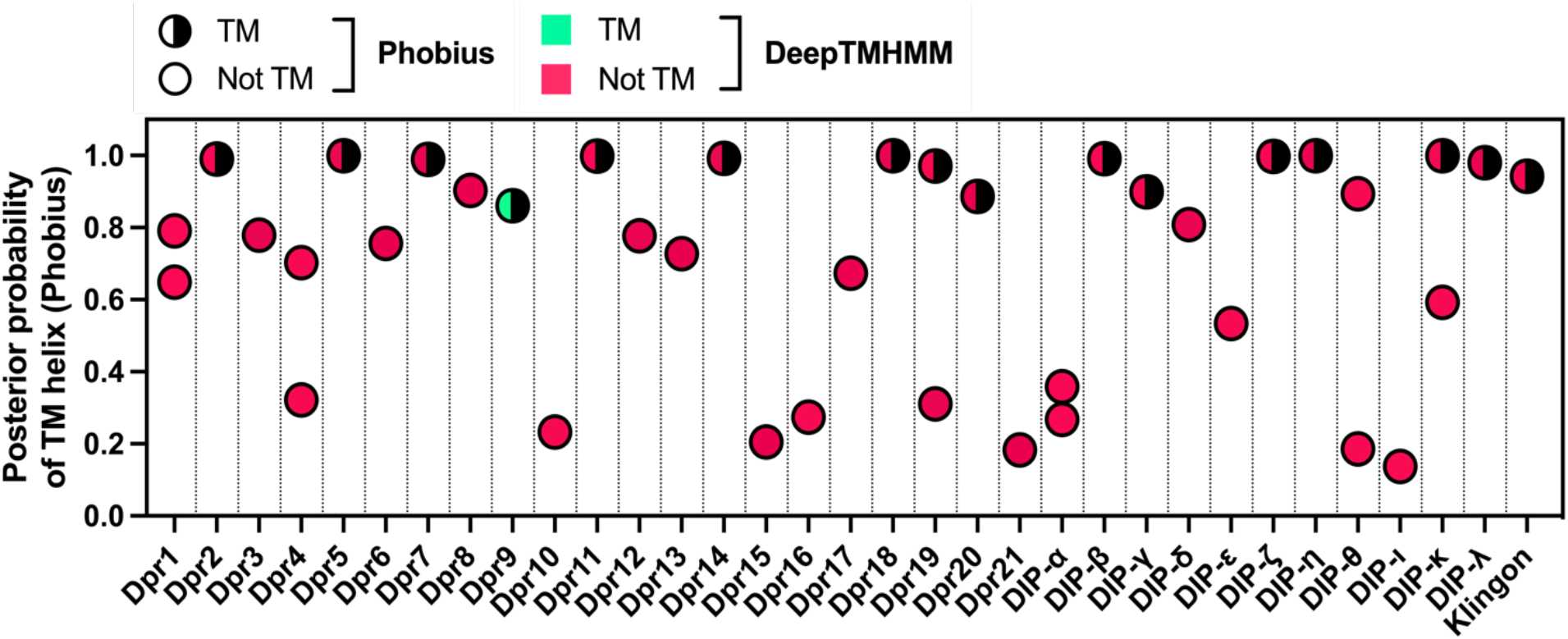
Transmembrane region prediction by Phobius and DeepTMHMM. Plotted are the highest values of posterior probability of a TM helix reported from Phobius. In cases where there were two regions with posterior probability of TM helix above 0.1, the highest values for both were plotted. TM, transmembrane. The highest values of posterior probability in Phobius do not always correlate with positive prediction for a given protein (Käll et al., 2007; for example, posterior probabilities of TM helix for Dpr8 or DIP-θ are higher than the one for Dpr9, but the former two were not predicted by Phobius to have a TM helix).

**Figure S2.**
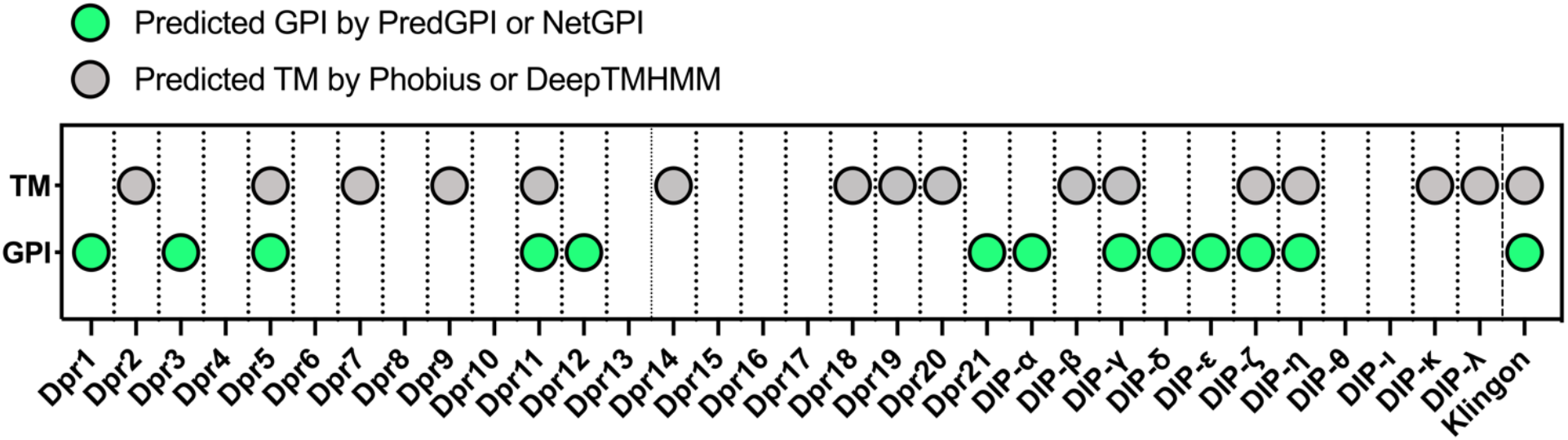
Summary of GPI anchor and TM helix predictions.

**Figure S3.**
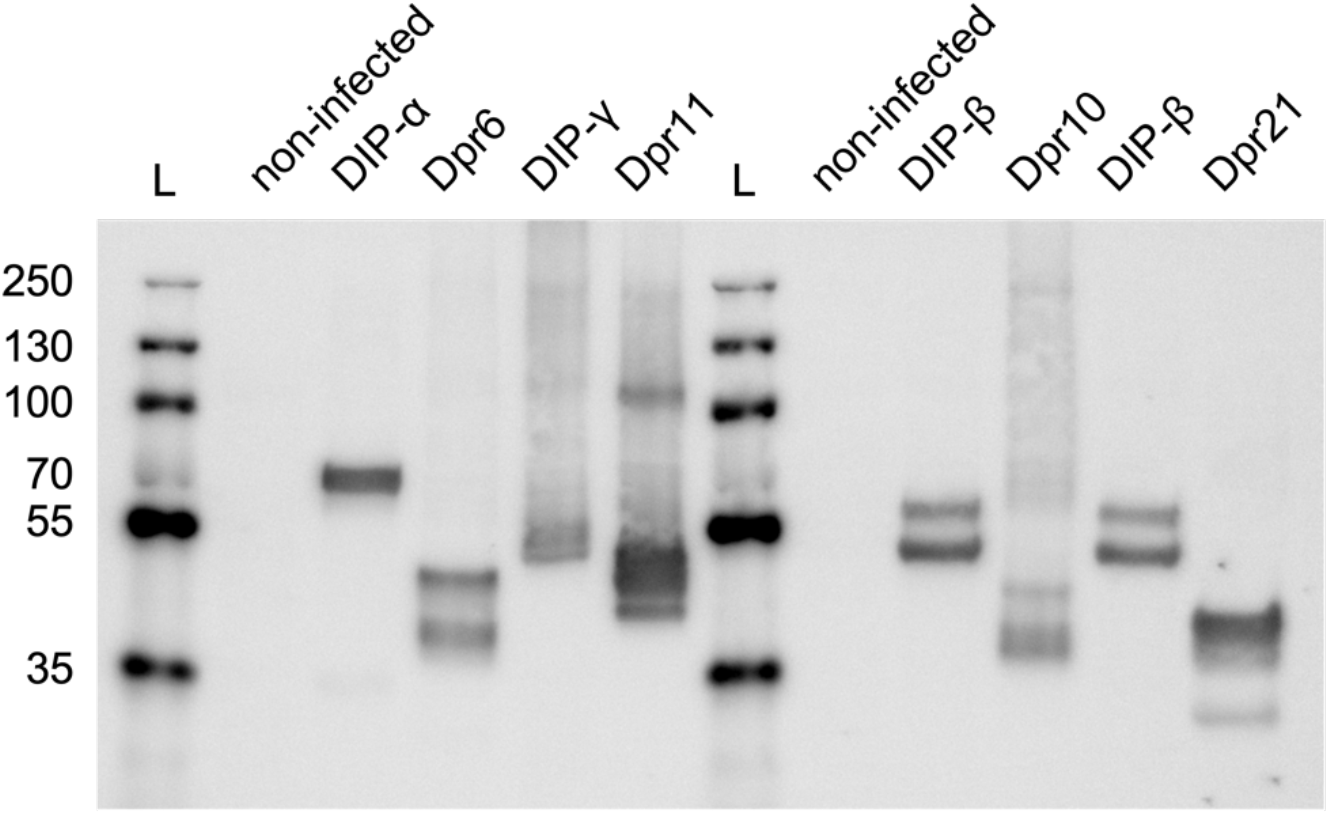
Expression of V5-tagged Dprs and DIPs in Sf9 cells used in cell aggregation assays. Sf9 cell pellets were solubilized and used in western blot with anti-V5-Alexa Fluor 647 antibody. L, molecular weight ladder.

**Figure S4:**
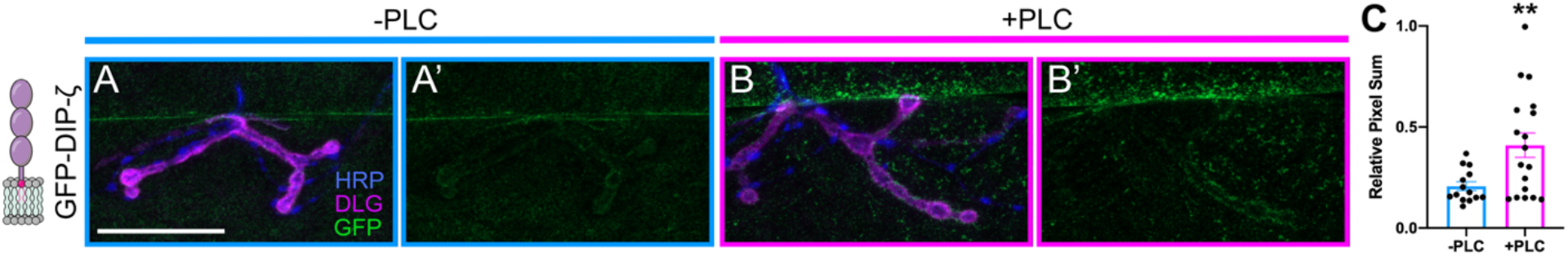
DIP-ζ aggregates in response to PLC treatment. (A-B’) Tagged DIP-ζ was expressed in muscles and dissected larvae were treated with PLC and compared to non-treated controls. Cartoon of tagged DIP-ζ is shown in left column. Surface localized DIP-ζ is shown in green, neuronal tissue in blue (HRP), and postsynaptic membrane in magenta (DLG). (A-B) PLC efficiently increased surface labeling of V5 on muscles expressing EGFP-DIP-ζ. Scale bar = 50 μm. (C) Quantification of experiments shown in A-B’.

